# Transcriptomic responses to hypoxia in endometrial and decidual stromal cells

**DOI:** 10.1101/2019.12.21.885657

**Authors:** Kalle T. Rytkönen, Taija Heinosalo, Mehrad Mahmoudian, Xinghong Ma, Antti Perheentupa, Laura L. Elo, Matti Poutanen, Günter P. Wagner

**Affiliations:** Institute of Biomedicine, Research Centre for Integrative Physiology and Pharmacology, University of Turku, Kiinamyllynkatu 10, 20014, Finland; Turku Bioscience Centre, University of Turku and Åbo Akademi University, Tykistökatu 6, 20520, Turku, Finland; Yale Systems Biology Institute, West Haven, Connecticut 06516, USA; Department of Ecology and Evolutionary Biology, Yale University, New Haven, CT 06511, USA; Department of Future Technologies, University of Turku, FI-20014 Turku, Finland; Department of Obstetrics and Gynecology, Turku University Hospital, Kiinamyllynkatu 4-8, 20521, Turku, Finland; Department of Obstetrics, Gynecology and Reproductive Sciences, Yale Medical School, New Haven 06510, USA; Department of Obstetrics and Gynecology, Wayne State University, Detroit, MI- 48201, USA

**Author notes:** Correspondence should be addresses to K T Rytkönen;. Address: Institute of Biomedicine, Research Centre for Integrative Physiology and Pharmacology, University of Turku, Kiinamyllynkatu 10, 20014, Finland / Turku Bioscience Centre, University of Turku and Åbo Akademi University, Tykistökatu 6, 20520, Turku, Finland.

**Keywords:** Endometrium, Oxygen, Hypoxia, Endometrial stromal fibroblasts, Decidual stromal cells, Transcription, Endometriosis, Promoter, JunD Proto-Oncogene, JUND, CCAAT Enhancer Binding Protein Delta, CEBPD

## Abstract

Human reproductive success depends on a properly decidualized uterine endometrium that allows implantation and the formation of the placenta. At the core of the decidualization process are endometrial stromal fibroblasts (ESF) that differentiate to decidual stromal cells (DSC). As variations in oxygen levels are functionally relevant in endometrium both upon menstruation and during placentation, we assessed the transcriptomic responses to hypoxia in ESF and DSC. In both cell types hypoxia upregulated genes in classical hypoxia pathways such as glycolysis and the epithelial mesenchymal transition. In DSC hypoxia restored an ESF like transcriptional state for a subset of transcription factors that are known targets of the progesterone receptor, suggesting that hypoxia partially interferes with progesterone signaling. In both cell types hypoxia modified transcription of several inflammatory transcription factors that are known regulators of decidualization, including decreased transcription of *STATs* and increased transcription of *CEBPs*. We observed that hypoxia upregulated genes had a significant overlap with genes previously detected to be upregulated in endometriotic stromal cells. Promoter analysis of the genes in this overlap suggested the hypoxia upregulated Jun/Fos and CEBP transcription factors as potential drivers of endometriosis-associated transcription. Using immunohistochemistry we observed increased expression of JUND and CEBPD in endometriosis lesions compared to healthy endometria. Overall the findings suggest that hypoxic stress establishes distinct transcriptional states in ESF and DSC, and that hypoxia influences the expression of genes that contribute to the core gene regulation of endometriotic stromal cells.

## Introduction

Human reproductive success depends on a properly differentiated (decidualized) uterine endometrium that allows implantation, the formation of the placenta and maintenance of the pregnancy (Gellersen and Brosens, 2014; Vinketova *et al.*, 2016). In humans, and other catarrhine primates, decidualization of endometrial stromal fibroblasts (ESF) to endometrial stromal cells (DSC) takes place spontaneously during every menstrual cycle. Decidualization involves substantial transcriptional and cellular remodeling, enabling implantation and placental development as well as menstruation-associated renewal of endometrium (Gellersen and Brosens, 2014). This process is triggered by autocrine and paracrine signaling pathways dependent on progesterone activated progesterone receptor (PGR) and cyclic adenosine monophosphate (cAMP) mediated activation of protein kinase A (PKA) (Pavličev *et al.*, 2017; Wu *et al.*, 2018). These, together with expression of transcription factors (TFs) including forkhead box protein O1 (FOXO1), homeobox (HOX), and signal transducer and activator of transcription (STAT) paralogs contribute to the regulatory programming necessary for decidualization (Gellersen and Brosens, 2014; Vinketova *et al.*, 2016).

The endometrium is exposed to hypoxic periods specifically upon menstruation as well as during placentation (Pringle *et al.*, 2010; Maybin and Critchley, 2015), but the associated transcriptional regulation remains poorly characterized. A recent study assessed the role of hypoxia in menstrual repair (Maybin *et al.*, 2018), but the difference in the hypoxia related transcriptional regulation between undifferentiated ESF and differentiated DSC has not been studied. Moreover, it was recently shown that DSC specific gene regulatory network involves factors that are part of the oxidative stress responses (Erkenbrack *et al.*, 2018), placing a specific interest on oxygen dependent gene regulation in the endometrium.

Importantly, endometrial responses to hypoxia are also relevant to several aspects of reproductive health. In endometriosis, endometrial cells grow outside of uterus in niches that are often more hypoxic than the highly vascularized uterus (Bishop, 1956; Bourdel *et al.*, 2007; Wu *et al.*, 2019). Endometriosis lesions grow faster in hypoxia (Lu *et al.*, 2014) and associated angiogenesis (Lu *et al.*, 2014) and hormone actions, including regulation of estrogen receptor (Wu *et al.*, 2012), are affected by hypoxia. Additionally, the abnormalities in pregnancy disorder preeclampsia are partly driven by hypoxia signaling (Tal, 2012).

Up to date no whole genome studies are available that describe the transcriptomic responses to hypoxia in the endometrial stromal cells. Here we characterize the transcriptomic responses to severe hypoxia (1% O_2_, 24h) in cultured ESF and DSC. We assess the hypoxia-regulated pathways by enrichment analysis and specifically focus on hypoxia regulated transcription factors. Further, we show that hypoxia upregulated genes have significant overlap with genes known to be upregulated in endometriosis, and guided by promoter analysis of transcription factor binding sites we select two transcription factors, JunD Proto-Oncogene (JUND) and CCAAT Enhancer Binding Protein Delta (CEBPD), for immunohistochemistry in endometriosis lesions and healthy endometria.

## Material and methods

### Cell Culture and hypoxia treatment

Human immortalized endometrial stromal fibroblasts (ESF) (T HESC, Mor lab, Yale University, corresponding to ATCC CRL-4003) were grown in normoxia (21% O_2_) in Dulbecco’s Modified Eagle’s medium (DMEM) (Sigma-Aldrich, D2906), supplemented with 10% charcoal stripped calf serum (Hyclone), 1% antibiotic/antimycotic (ABAM; Gibco), 1nM sodium pyruvate (Gibco), 0.1% insulin-transferrin-selenium (ITS premix, BD Biosciences), and 0.12% sodium bicarbonate. To generate DSC, ESFs were decidualized by adding of 0.5 mM 8-bromoadenosine 3′,5′-cyclic monophosphate (8-Br-cAMP) (Sigma) and 0.5 μM of the synthetic progestin medroxyprogesterone acetate (MPA) in DMEM supplemented with 2% charcoal-stripped calf - serum.

For ESF hypoxia exposure was conducted for 24 hours using ProOx C21 nitrogen-induced hypoxia system (BioSpherix, Red Field, NY) at 1% O_2_, 5% CO_2_ and compared to normoxic ESF from the same cell batch. For DSC, ESF were first decidualized for 36 hours and then similarly exposed to hypoxia for 24 hours (total decidualization time including the time under hypoxia = 60 hours) and compared to normoxic DCS from the same cell batch that were decidualized for two days. As hypoxia slows down cellular processes we defined that normoxic decidualization of 36 hours followed by 24 hours decidualization in hypoxia represents a reasonable approximation to be compared to normoxic decidualization of two days. Each of the four sample groups had two biological replicates.

### RNA-seq, differential transcription and visualization

Total RNA was extracted with RNeasy Mini or Midi RNA-extraction kits (QIAGEN) followed by on-column DNase I treatment. Total RNA quality was assayed with a Bioanalyzer 2100 (Agilent) and 500 ng of RNA samples were sequenced with Illumina Genome Analyzer II platform. For each sample at least 30 million reads were acquired and quality parameters were checked with FastQC (http://www.bioinformatics.babraham.ac.uk/projects/fastqc). Sequencing data are available in NCBI Gene Expression Omnibus (GEO; https://www.ncbi.nlm.nih.gov/geo/) under accession numbers GSE111570 (GSM3034449, GSM3034450, GSM3034451 and GSM3034452) and GSE63733 (GSM1556296, GSM1556297, GSM1556298, GSM1556299). All data is also available from authors upon request.

Sequence reads were mapped to the GRCh37 human reference genome using Tophat2 (Trapnell *et al.*, 2009) and the gene counts were calculated using HTSeq (Anders *et al.*, 2015) according to Ensembl annotation (GRCh37.69) and normalized as transcripts per million (TPM) (Wagner *et al.*, 2013). Differential transcription was analyzed with edgeR using upper quartile normalization (Robinson *et al.*, 2010) and the following cut-offs: false discovery rate (FDR) < 0.01, absolute fold-change (FC) > 2.0 and TPM > 2. The principal component analysis (PCA) of the four conditions after edgeR upper quartile normalization indicated that the two biological replicates in each condition grouped tightly together (Supplementary Figure 1).

For visualization, gene heatmaps were produced from averages of the absolute TPM values using pheatmap_1.0 in R 3.5. Hierarchical clustering of the genes was performed using Euclidean distance and the complete linkage method. TF set was constructed by combining genes in Ingenuity Pathway Analysis (IPA, Qiagen, www.qiagen.com/ingenuity) categories “transcription regulator” and “ligand-dependent nuclear receptor”. Hypoxia regulated transcription factor subsets were intersected with PGR targets that were previously detected using siRNA by Demayo lab (we filtered the originally reported siPGR set using FC > 2) (Mazur *et al.*, 2015). A circos plot was produced in METASCAPE (metascape.org).

### Pathway analysis

For gene ontology (GO) and gene set enrichment analysis and heatmaps, gene lists of differentially transcribed genes (FDR < 0.01, FC > 2.0, TPM > 2) were used as input for METASCAPE, and available pathway databases (GO Biological Processes, Reactome Gene Sets, Canonical Pathways, Biocarta Gene Sets, KEGG Pathway and Hallmark Gene Sets) were selected for analysis. We used “IPA Canonical Pathways” tool to visualize enriched pathways.

### Endometriosis transcriptome data and statistical tests of the overlaps

In order to test the significance of our cell culture results in a clinically relevant hypoxic niche we investigated the overlap of hypoxia differentially transcribed genes with most relevant available endometriosis data from literature and databases. Endometriosis data sets included a stromal dataset (FACS isolated with CD10 antibody) of differentially expressed genes between endometriosis lesions versus control healthy endometrium (Rekker *et al.*, 2017, Supplementary Table 1); a heterogeneous tissue dataset of differentially expressed genes in the endometrium of endometriosis patients versus healthy controls (Tamaresis *et al.*, 2014, Supplementary Table 5, endometriosis and abnormal) and endometriosis related genes listed in the DisGeNET v5.0-database (http://www.disgenet.org/) (Piñero *et al.*, 2017). For the first two sets, lists of endometriosis up- and downregulated genes were analyzed separately, whereas for the database one list of all endometriosis-regulated genes was used. Each list was filtered to contain only genes that were expressed in our ESF and DSC cell cultures (TPM > 2). To study the overlaps of the endometriosis differentially transcribed genes and the hypoxia differentially transcribed genes we organized the data using varhandle 2.0.3 and constructed contingency tables in R 3.6.0, conducted Fisher’s exact test (fisher.test) to determine the significance of enrichment and presented the results using pheatmap 1.0.12. Effect size was calculated as odds ratio, which was provided by the fisher.test. The genes of the two most significant overlaps were further analyzed with pathway analysis and transcription factor binding site motif analysis.

### Transcription factor binding site (TFBS) analysis

We tested the enrichment of TFBS in the promoters of hypoxia and endometriosis upregulated genes using GeneXplain 4.0 (http://genexplain.com). The endometriosis upregulated subsection of all the hypoxia upregulated genes was set as foreground (yes set) and all hypoxia upregulated genes as the background/control (no set) and “Search for enriched TFBSs (genes)” function was run with default settings (promoter: −1000 to +100). Two TFBS databases were searched: prediction based TRANSFAC_public_vertebrates and ChIP-seq based GTRD_moderate. Both result lists were ranked with “Site FDR” to inspect the most significantly enriched TFBS motifs. Top 20 enriched motifs were manually examined for corresponding TFs homologs among the top ESF and DSC hypoxia upregulated TFs. As TFBS motif names and TF names do not constitute one-to-one correspondence, corresponding homologs (and synonyms) and were manually defined using https://www.genecards.org/.

### Immunohistochemistry

Patient samples were collected and processed as described previously (Heinosalo *et al.*, 2018). Briefly, the Joint Ethics Committee of Turku University and Turku University Hospital approved collections and all study subjects provided written informed consent. Samples of endometriosis (deep infiltrating lesions: sacrouterine ligament and bladder) and eutopic endometrial biopsies were collected from endometriosis patients, and as a control group, endometrial biopsies from healthy, endometriosis-free women undergoing laparoscopic tubal ligation were collected. Tissue samples were fixed in formalin and embedded in paraffin for histological analysis. Antigen retrieval of hydrated 5 μm tick sections was performed in 10 mM sodium citrate buffer (pH 6.0), followed by immunohistochemistry with primary antibodies against JUND (mouse monoclonal, Jun D Antibody (D-9), sc-271938, Santa Cruz Biotechnology, Santa Cruz, CAUSA) with 1:1500 dilution and CEBPD (rabbit polyclonal Anti-CEBP Delta antibody ab65081, Abcam, Cambridge, MA, USA) with 1:250 dilution. Three samples in each sample group (endometriosis lesion, patient endometrium, healthy control endometrium) were stained. Sections were scanned for analyses with the panoramic 250 Flash series digital slide scanner (3DHISTECH, Hungary).

## Results

### Global transcriptomic effects of hypoxia

We performed global RNA-seq from hypoxia treated (1% O_2_, 24h) immortalized human endometrial stromal fibroblasts ESF (ATCC CRL-4003) and 2 day decidualized (MPA, 8-br-cAMP) endometrial stromal cells (DSC) and compared these to corresponding normoxic conditions. Hypoxia-related upregulation of transcription (FDR < 0.01, FC > 2 and TPM > 2 in hypoxic condition) involved more shared genes between ESF and DSC compared to hypoxia-related downregulation of transcription (FDR < 0.01, FC < −2 and TPM > 2 in normoxic condition) (Fig. 1A). Specifically, 36% (728) of the genes upregulated were shared in both ESF and DSC whereas 628 and 643 genes, respectively, were upregulated in a cell type specific manner. In contrast, 69% of the genes downregulated in DSC (1304) were DSC specific (901).

**Figure 1.**
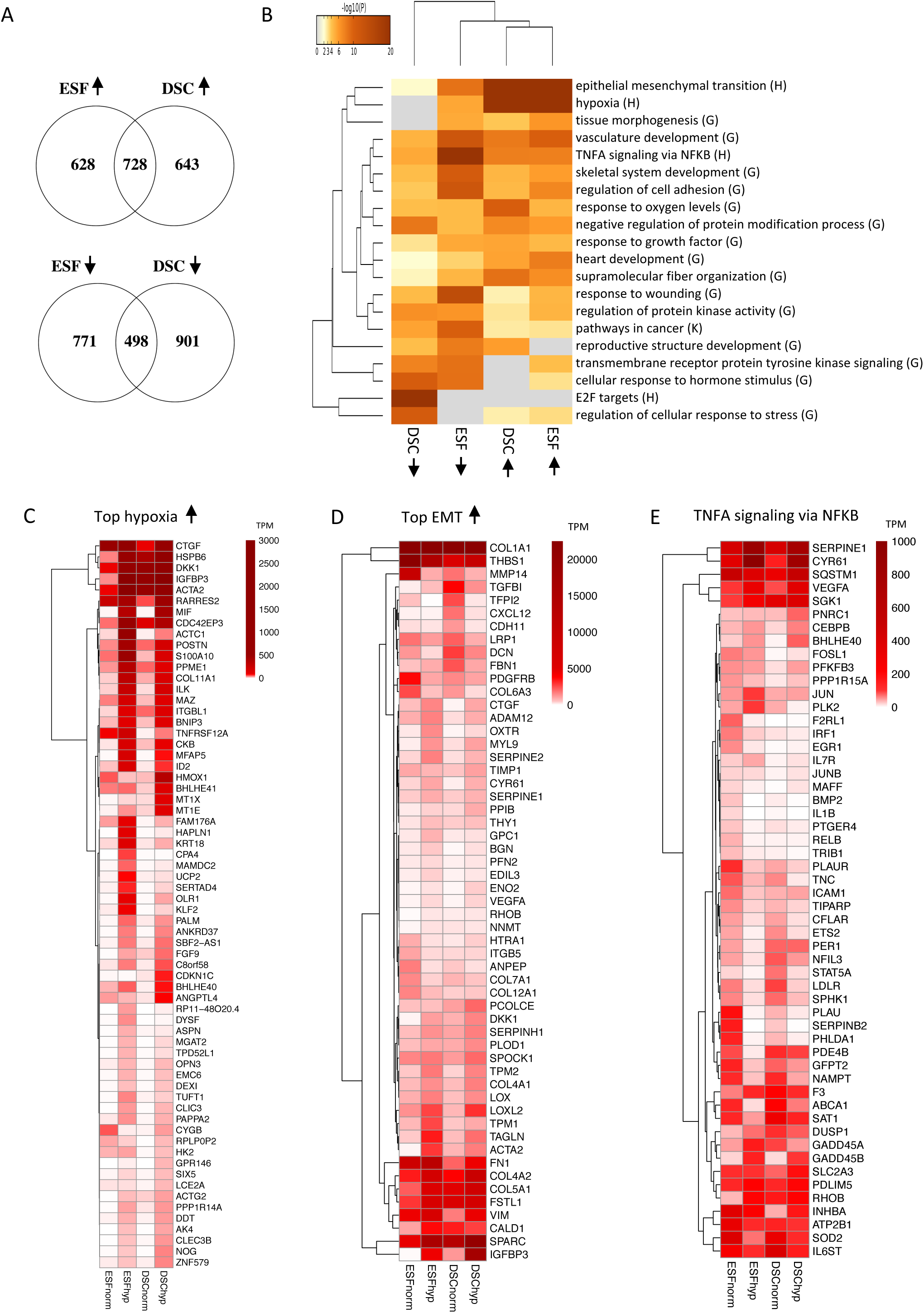
Hypoxia regulated gene pathways in endometrial stromal fibroblasts (ESF) and endometrial stromal cells (DSC). A) Differentially transcribed genes in hypoxia (1% O_2_, 5% CO_2_, 24 hours, FC > 2, FDR < 0.01, TPM > 2). Arrows up mark upregulated genes and arrows down mark downregulated genes. B) Hierarchical clustering of the functional enrichment categories from METASCAPE (*P* values, default; see Supplemental Table S1) using the genes with significantly increased and decreased transcription as an input. The functional categories are from GO Biological Processes (G), Hallmark gene sets (H) and KEGG (K). Arrows mark the upregulated or downregulated genes in the four conditions. C) Top 40 genes with increased transcription in hypoxic conditions in ESF and DSC (norm = normoxia, hyp = hypoxia), presented using hierarchical clustering (Euclidean, complete) of averages of absolute TPM values (> 20 TPM). D) “Epithelial mesenchymal transition” (EMT) term associated hypoxia upregulated and highly transcribed genes (> 200 TPM). E) All “TNFA signaling via NFKB” term associated genes differentially transcribed under hypoxia (> 20 TPM). TNFA, Tumor Necrosis Factor A; NFKB, Nuclear Factor Kappa B. See Supplementary Table 1 for the full lists of hypoxia regulated genes, the gene lists of detected enrichment categories and TPM values used for the figures.

### Functional pathways affected by hypoxia

Pathway enrichment analysis revealed a clustering pattern of the functional terms that is concordant with the above described gene proportions. In both cell types the most significant cellular functions predicted to be upregulated by hypoxia (“hypoxia” and “epithelial mesenchymal transition”) were clustered together (Fig. 1B). On the other hand hypoxia downregulated genes in ESF and DSC formed distinct enrichments, in ESF the most significant terms being “TNFA signaling via NFKB” and “response to wounding” whereas in DSC these were “E2F targets” (involved in cell cycle regulation) and “regulation of cellular response to stress” that were not detected in ESF.

As expected, in both cell types hypoxia upregulated genes included canonical glycolysis pathway (Supplementary Figure 2) and other known target genes of hypoxia inducible factor 1 (*HIF-1alpha*) (Fig. 1C). These genes, which also shared the highest hypoxic increases in both conditions, included insulin-like growth factor binding protein 3 (*IGFBP3*), macrophage migration inhibitory factor (*MIF*) and metallothionein genes (*MT1E, MT1X*). *MIF* is a cytokine potentially regulating both inflammatory status and angiogenesis (Hahne *et al.*, 2018), and *MT1*s are involved in the protective responses to oxidative stress (Xue *et al.*, 2012). Upon hypoxia in DSC, of the classic decidualization markers, transcription of prolactin remained on the same level as in normoxia (TPM 7.9 to 5.4), whereas *IGFBP1* was markedly downregulated (TPM 17 to 0.4) suggesting that hypoxia interferes especially with metabolic aspects of decidualization. Most notably, in hypoxia the TPM for *IGFBP3*, a paralog of *IGFBP1*, increased from 38 to 4382 TPM in ESF, and from 1216 to 16865 TPM in DSC. IGFBPs bind to and modulate insulin-like growth factors (IGFs) and may affect glucose uptake and proliferation of the cells. Also LDHA was highly transcribed and 8.6-fold upregulated in ESF (to TPM 1645) and 5.3-fold upregulated in DSC (to TPM 1381) (Supplementary Table 1A) confirming the extensive hypoxic metabolism together with glycolysis pathway.

In our study, the top hypoxia upregulated genes with high transcription levels included several extracellular matrix components and their regulators, including several collagens (1A1, 6A3, 7A1, 12A1, 4A1, 4A2, 5A1), fibronectin, vimentin, proteases and Lysyl oxidases that are involved in EMT (Fig. 1D) and that are relevant markers of mesenchymal like state in ESF (Yu *et al.*, 2016; Owusu-Akyaw *et al.*, 2019). Thus, knowing that these genes are central for both EMT and the opposite process mesenchymal epithelial transition (MET), these results suggest that hypoxia may repress mesenchymal epithelial transition (MET)-like extracellular modifications typical for early decidualization.

In ESF, hypoxia repressed genes connected to inflammation, under the terms such as “TNFA signaling via NFKB” and “response to wounding” (Fig. 1B). This repression included the downregulation of pro-inflammatory cytokines such as *IL1B* and *BMP2*, and TFs *RELB, F2RL1* and *IRF1* (Fig. 1E). These results suggest that hypoxia modifies the pro-inflammatory signaling characteristic for early decidualization (Salker *et al.*, 2012; Rytkönen *et al.*, 2019). In DSC, hypoxia downregulated genes in the “E2F targets” term that includes several regulators of cell cycle, chromosome segregation and nuclear division (Supplementary Table 1B). The DSC downregulated genes within the term “regulation of cellular response to stress” include multitude of genes in intracellular protein kinase signaling such as MAPK and JNK cascades (Supplementary Table 1B).

### Hypoxia regulated transcription factors

The TFs upregulated in response to hypoxia included several common mediators of hypoxia-induced gene repression. Of the TFs upregulated in ESF by hypoxia, 60% (66/110) were also upregulated in DSC, whereas of the downregulated TFs only 37% (37/100) were shared (Fig. 2A). In both cell types hypoxia upregulated transcriptional repressors, such as Inhibitor Of DNA Binding (*ID*) genes that repress the expression of other TFs (Fig. 2B).

**Figure 2.**
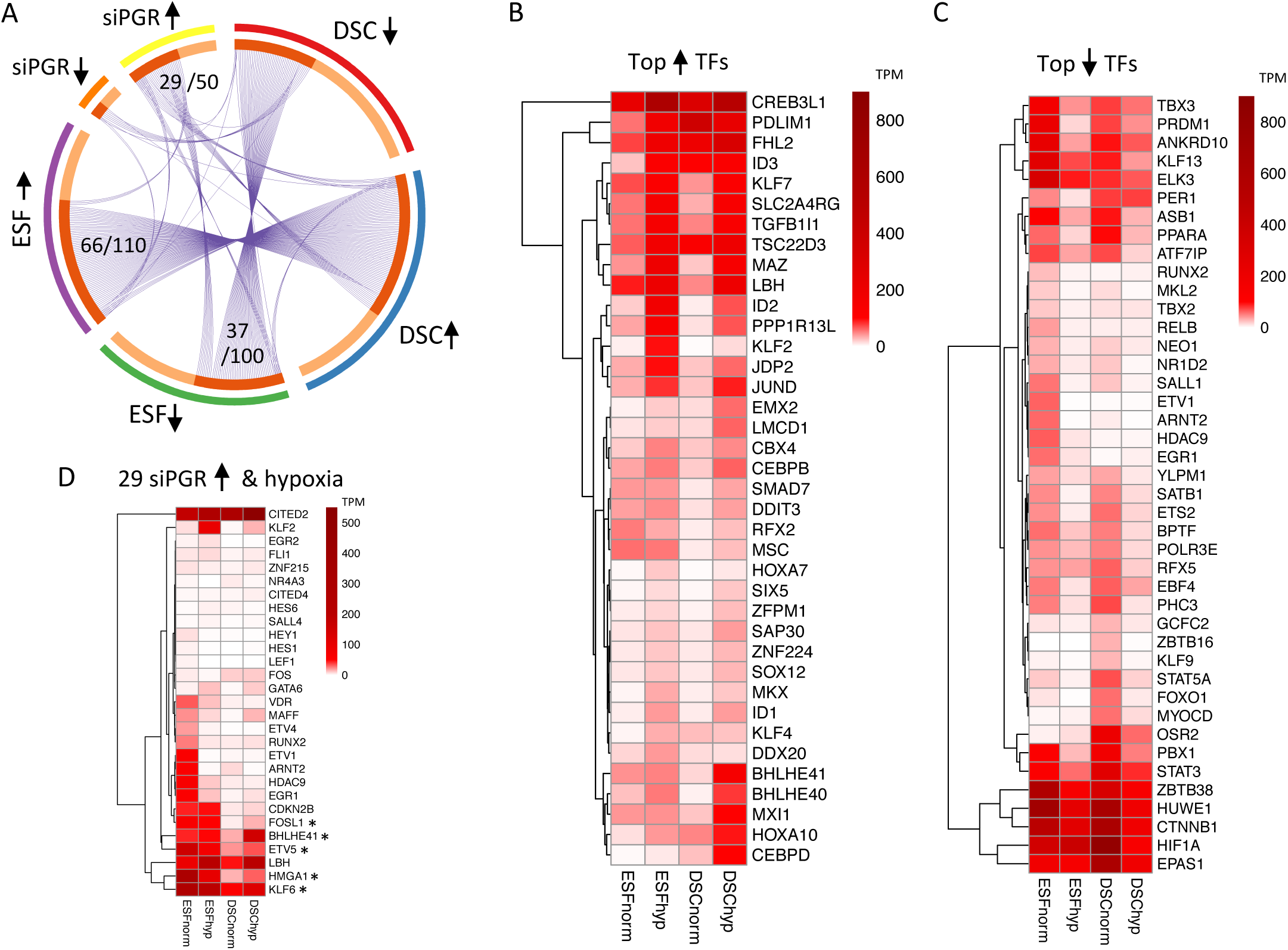
Top hypoxia regulated transcription factors (TF) in endometrial stromal fibroblasts (ESF) and endometrial stromal cells (DSC). A) Differentially transcribed TFs in hypoxia (1% O_2_, 5% CO_2_, 24 hours, FC > 2, FDR < 0.01, TPM > 2) and TFs regulated by decidualization regulator progesterone receptor (PGR) PGR siRNA (Mazur *et al.*, 2015) (FC > 2). Arrows up mark upregulated genes and arrows down mark downregulated genes. Top 25 TFs with increased (B) or decreased (C) transcription in ESF and DSC under hypoxia (norm = normoxia, hyp = hypoxia) presented using hierarchical clustering (Euclidean, complete) of averages of absolute TPM values (> 20 TPM). D) 29 Hypoxia regulated (ESF and/or DSC) and PGR downregulated (siRNA upregulated) TFs presented using hierarchical clustering (Euclidean, complete) of averages of absolute TPM values (> 20 TPM). * Notes a subset of highly transcribed and PGR downregulated TFs for which hypoxia partly restores the ESF like transcriptional state (*FOSL1, KLF6, BHLHE41, HMGA1, ETV5*). *FOSL1*, FOS Like 1, AP-1 Transcription Factor Subunit; *KLF6*, Kruppel Like Factor 6; *BHLHE41*, Basic Helix-Loop-Helix Family Member E41 (DEC2); *HMGA1*, High Mobility Group AT-Hook 1; *ETV5*, E26 transformation-specific Variant 5. See Supplementary Table 2 for the lists of hypoxia and PGR regulated TFs and the lists of the specific intersections with corresponding TPM values.

The master hypoxic regulators (*HIF-1alpha* and *EPAS1*, a.k.a. *HIF-2alpha*) were highly transcribed in all conditions, but were downregulated in DSC by hypoxia 5- and 3-fold, respectively (Fig. 2C). Notably, in DSC, *BHLHE40* and *BHLHE41* were highly upregulated by hypoxia, 17-fold and 11-fold respectively (Fig. 2B). These repress TFs regulating circadian rhythm, such as PER and CLOCK. In line with this, we observed hypoxic downregulation of *PER1/3* in ESF and *CLOCK* in DSC (Supplementary Table 1A).

The effects of hypoxia on core regulators of decidualization were multifaceted. In both cell types, and particularly in DSC, *HOXA10, HOXA11, CEBPB* and *CEBPD* were upregulated by hypoxia (Fig. 2B, Supplementary Table 1A), whereas *FOXO1, STAT3* and *STAT5A* were downregulated (Fig. 2C). Of these especially CEBPs and STATs are involved in the initial pro-inflammatory decidualization phase (Wang *et al.*, 2012; Rytkönen *et al.*, 2019).

Other notable upregulated groups of TFs included members of Jun/Fos family (JUND and JDP2) and several Kruppel-like factors such as KLF2, KLF4 and KLF7, of which KLF2 was 18-fold upregulated in ESF. KFL2 is a known negative regulator of NFKB pathway (Jha and Das, 2017), and potentially participates in the of the observed ESF specific NFKB pathway repression.

### Hypoxia interferes with Progesterone Receptor regulated TF networks

In both cell types hypoxia downregulated several genes that were under the GO term “cellular response to hormone stimulus” (Fig. 1B). Signaling via progesterone receptor (PGR) is known to be a main driver of decidualization, and it regulates e.g. the expression of *FOXO1, HOXO10, CEBP*s, and *STATs*. We, thus, investigated the effect of hypoxia specifically in PGR regulated TFs by intersecting ChIP-seq detected and PGR regulated (>2-fold) TFs from a previous study (Mazur *et al.*, 2015) with the hypoxia regulated TFs. The data revealed that 8/18 PGR upregulated and 29/50 PGR downregulated TFs were influenced by hypoxia (Fig. 2A). This suggests that primarily TF networks downregulated by PGR are modified by hypoxia. Notably, of the 29 TFs downregulated by PGR, 13 showed similar downregulation by hypoxia in ESF, while 15 were upregulated in DSC by hypoxia (Fig. 2A, D). The data also suggest that in DSC hypoxia partly restores the ESF like transcriptional state for a subset of highly transcribed and PGR downregulated TFs (Fig. 2D). Overall these observations suggest that hypoxia, at least partly, reverses PGR dependent transcriptional changes necessary for early decidualization.

### Relationship between gene sets upregulated by hypoxia and in endometriosis

In order to explore the significance of our cell culture results for a clinically relevant hypoxic niche, we investigated the overlap between the genes differentially transcribed in hypoxia with the data available on endometriosis using Fisher’s exact test (Fig. 3A). The overlap with the highest significance was observed between the ESF hypoxia upregulated genes and the endometriosis upregulated genes (p = 5.8E-27, 3.90-fold) from a dataset comparing isolated endometrial stromal cells from endometriosis lesions to healthy control endometrium (Rekker *et al.*, 2017). In this dataset, also the overlap with DSC hypoxia upregulated genes with endometriosis upregulated genes was highly significant (p = 6.0E-19, 3.25-fold). This extensive overlap of hypoxia upregulated (ESF and DSC) genes of the endometriosis upregulated genes 39% (157/402, TPM > 2 in our cell culture) suggests that gene regulatory programs responding to hypoxic conditions contribute to the core gene regulation in the stroma of endometriosis lesions.

**Figure 3.**
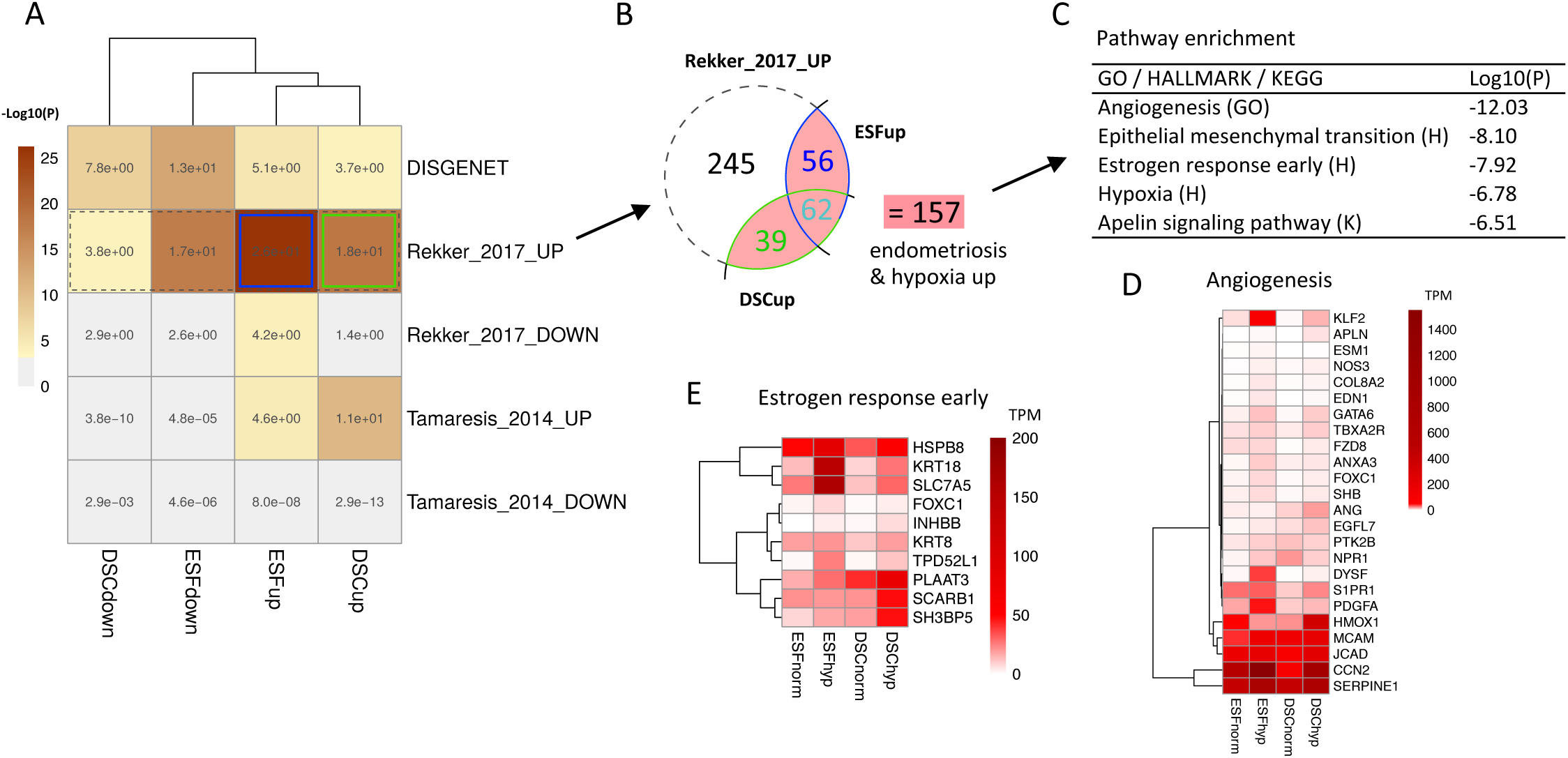
Overlaps of hypoxia regulated and endometriosis regulated genes. (A) Significance of the overlaps of hypoxia regulated genes and three representative endometriosis datasets determined using Fishers exact test. Overlaps between hypoxia upregulated genes with stroma specific endometriosis upregulated genes (Rekker *et al.*, 2017) were highly significant (blue and green boxes). Other analyzed datasets were a whole tissue (cell type heterogeneous) endometrium dataset of endometriosis patients versus healthy controls (Tamaresis *et al.*, 2014) and endometriosis associated genes from DisGeNET v5.0 database. (B) Venn diagram showing the number of genes in (Rekker *et al.*, 2017) endometriosis upregulated genes (dashed line) with overlapping ESF and DSC hypoxia upregulated genes (unique green and blue, respectively). Total of 157 genes that were hypoxia and endometriosis regulated were then used as an input for pathway enrichment analysis (C) of which the top enrichment categories are displayed. (D) Heatmap of genes in the “Angiogenesis” enrichment term. (E) Heatmap of genes in the “Estrogen response early” enrichment term. The functional categories are from GO Biological Processes (GO), Hallmark gene sets (H) and KEGG (K). See Supplementary Table 3 for the original and overlapping gene lists, enrichment gene lists and corresponding TPM values used for the heatmaps.

The overlaps with endometriosis datasets that were not stromal cell specific were not significant or less overlapping compared to the stromal cell specific data (Fig. 3A). For a heterogeneous tissue dataset of differentially expressed genes in the endometrium of endometriosis patients versus healthy controls (Tamaresis *et al.*, 2014), the overlaps of hypoxia and endometriosis upregulated genes were moderately significant (p = 2.5E-11 – 2.2E-5, 2.66-fold – 1.92-fold). Further, the endometriosis related genes in DisGeNET v5.0 database (without the direction of regulation) had moderately significant overlaps (p = 2.2E-13 – 2.0E-4, 2.58-fold – 1.64-fold) in both hypoxia up- and downregulated genes.

We then used the detected stromal cell specific 157 genes that were hypoxia and endometriosis upregulated as an input for pathways analysis (Fig. 3B). The most enriched categories included “angiogenesis”, “epithelial mesenchymal transition” and “estrogen response early” (Fig. 3C). The hypoxia and endometriosis regulated “angiogenesis” genes included several highly transcribed and secreted ECM and adhesion molecules such as CCN2 (Cellular Communication Network Factor 2), Serpine1, MCAM (Melanoma Cell Adhesion Molecule), JCAD (Junctional Cadherin 5 Associated) (Fig. 3D). Additionally, the known core endometriosis TF GATA6 was hypoxia upregulated and the previously mentioned KLF2 (Kruppel Like Factor 2) was substantially upregulated in hypoxic ESF (Fig. 3D). Of the hypoxia and estrogen regulated genes KRT8 and KRT18 are keratins that dimerize with each other to form intermediate filaments, and SCL7A5 (Solute Carrier Family 7 Member 5, or LAT1) is an amino-acid transporter (Fig. 3E).

### Promoter transcription factor binding site (TFBS) analysis of hypoxia and endometriosis upregulated genes reveals enrichment for Jun/Fos and CEBP motifs

We next analyzed the promoters of these 157 hypoxia and endometriosis upregulated genes (Fig. 3B) to find TFs that are both hypoxia upregulated and potential drivers of endometriosis. Using GeneXplain 4.0 we searched two databases, prediction based TRANSFAC and ChIP-seq based GTRD, for TFBS motifs enriched in the promoters (−1000 to +100 of TSS) of these 157 genes (Table 1). Then we manually examined the top 20 ranked (site FDR) motifs for corresponding TFs homologs among the highly hypoxia expressed and upregulated TFs (from Fig. 2B). We discovered that in both TRANSFAC and GTRD databases there were corresponding homolog motifs for the hypoxia upregulated Jun/Fos family members JDP2 and JUND and CEBP family members CEBPB and CEBPD. Of these JUND and CEBPD were selected for protein level validation in endometriosis biopsies due to their higher hypoxic upregulation compared to JDP2 and CEBPB (Fig. 2B) in ESF (JUND 3.2-fold, CEBPD 5.0-fold) and in DSC (JUND 4.8-fold, CEBPD 5.2-fold).

**Table 1.** Transcription factor binding site (TFBS) enrichment analysis in the promoters of hypoxia (ESF and DSC) and endometriosis upregulated genes. Both prediction based (TRANSFAC) and ChIP-seq based (GTRD) databases were searched for TFBS enrichment using the 157 hypoxia and endometriosis upregulated genes as foreground and all hypoxia upregulated genes as a background. Top enriched TFBS motifs are displayed by “Site FDR” rank and “Hypoxia up” column displays the presence of a TF homologs corresponding to the TFBS motifs among top hypoxia upregulated TFs (from Fig. 2B). Endometriosis upregulated genes are from (Rekker *et al.*, 2017). See Supplementary Table 4 for the input gene lists and full TFBS enrichment lists.

| TRANSFAC | Site FDR | Hypoxia up | GTRD | Site FDR | Hypoxia up |
| --- | --- | --- | --- | --- | --- |
| OCT1_04 | 4,1E-49 | na | MEF-2A.1 | 7,5E-17 | na |
| TATA_01 | 4,1E-49 | na | B-ATF.1 | 7,5E-17 | na |
| MEF2_03 | 3,1E-38 | na | MEF-2C.1 | 3,5E-15 | na |
| RSRFC4_01 | 1,7E-32 | na | CEBPalpha.1 | 7,5E-13 | CEBPB, CEBPD |
| FOXJ2_02 | 1,5E-28 | na | TBP.1 | 5,7E-10 | na |
| MEF2_01 | 1,7E-26 | na | STAT6.1 | 1,1E-06 | STAT4 |
| PBX1_01 | 8,4E-26 | na | SOX-2.1 | 1,1E-06 | SOX12 |
| MEF2_02 | 2,8E-25 | na | GATA-2.1 | 1,1E-06 | na |
| GATA2_01 | 2,4E-24 | na | c-Jun.1 | 2,9E-06 | JDP2, JUND |
| HNF1_01 | 5,6E-23 | na | TCF-7L1.1 | 6,7E-06 | na |
| CDP_01 | 3,3E-21 | na | SMAD1.1 | 9,6E-06 | SMAD7 |
| AP1_01 | 1,5E-20 | JDP2, JUND | CEBPdelta.1 | 1,1E-05 | CEBPB, CEBPD |
| BRN2_01 | 1,3E-19 | na | FOXP2.1 | 1,4E-05 | FOXP4 |
| CEBP_Q2 | 2,0E-19 | CEBPB, CEBPD | PBX-1.1 | 1,6E-05 | na |
| AREB6_04 | 7,0E-19 | na | IRF-4.1 | 2,1E-05 | na |
| EVI1_05 | 1,4E-17 | na | PU1.1 | 2,3E-05 | na |
| PAX4_04 | 1,6E-17 | na | STAT5A.1 | 7,9E-05 | STAT4 |
| OCT1_01 | 4,0E-17 | na | STAT1.1 | 2,8E-04 | STAT4 |
| EN1_01 | 6,3E-17 | na | GATA-3.1 | 3,0E-04 | na |
| MEF2_04 | 2,0E-16 | na | PPARgamma | 3,2E-04 | na |

The top motifs in the TFBS analysis that did not have corresponding hypoxia upregulated TF homolog in our experiment included motifs for TFs such as MEF2 (Myocyte Enhancer Factor 2), B-ATF (Basic Leucine Zipper ATF-Like TF), OCT1 (POU2F1, POU Class 2 Homeobox 1) and TBP/TATA (TATA binding protein) (Table 1). Of the homologs of these, MEF2C was 2.6-fold downregulated in ESF, whereas the others were not hypoxia regulated. Notably, B-ATF is AP-1/ATF family TF that dimerizes with JUN paralogs, but BATF genes (*BATF1, BATF2, BATF3*) were not expressed in our cell cultures (TPM< 2).

### Elevated immunohistochemical staining of JUND and CEBPD in deep endometriosis compared to endometrial biopsies

Immunohistochemical staining of JUND and CEBPD was conducted in deep infiltrating endometriosis lesions and compared to endometrial biopsies (patients and healthy controls) (Fig. 4). The stromal staining of JUND was considerable stronger in endometriosis compared to control endometrium, however, in both roughly only half of the cells were stained. Strong epithelial staining for JUND was present in all epithelial cells of the lesion whereas in control endometrium epithelial cell staining was weak. For CEBPD we detected staining in most of the stromal and epithelial cells, and both stromal and epithelial cells had stronger staining in the endometriosis lesions compared to control endometria.

**Figure 4.**
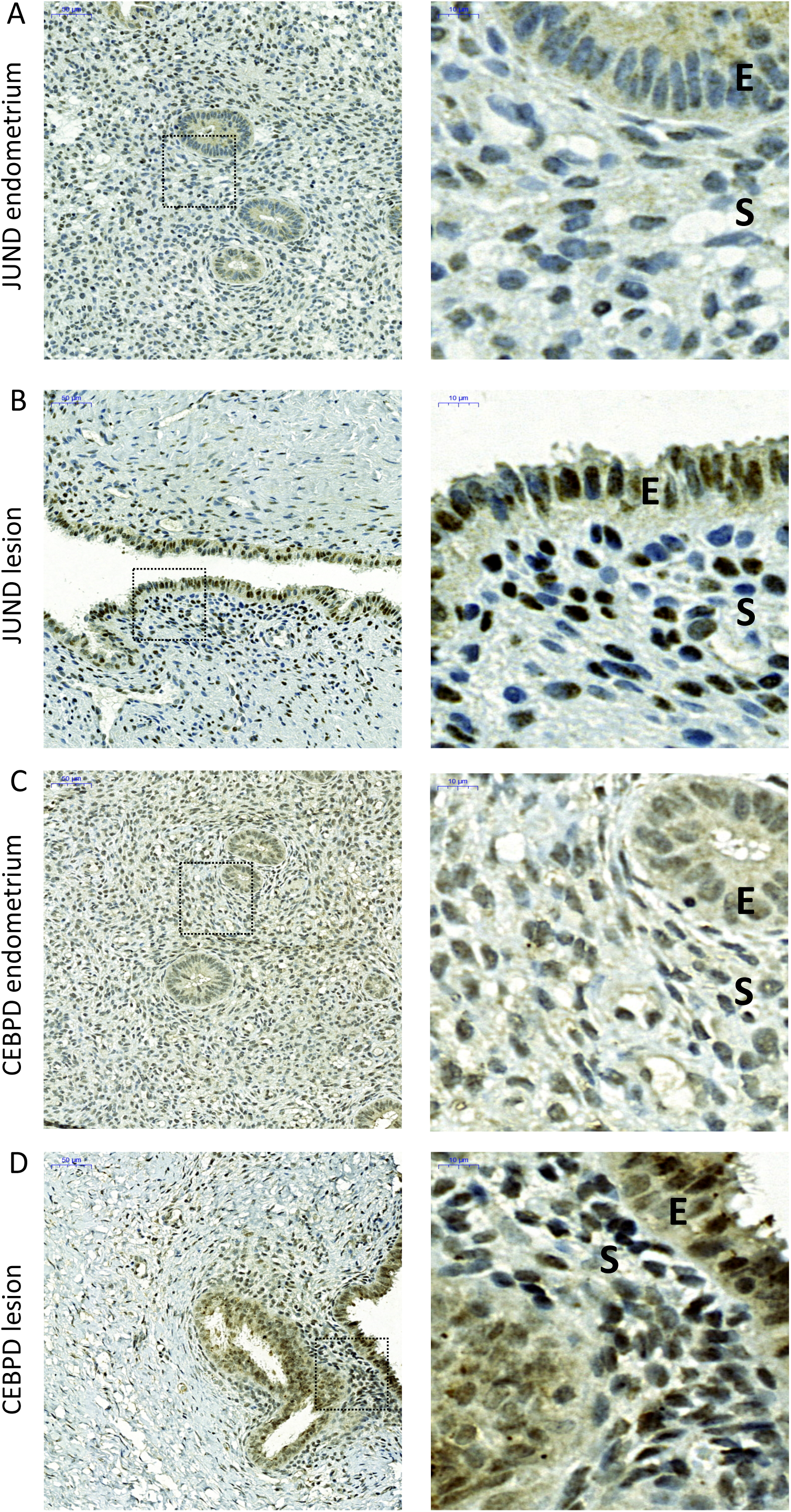
Immunohistochemical staining of JunD Proto-Oncogene (JUND) and CCAAT Enhancer Binding Protein Delta (CEBPD) in deep infiltrating endometriosis lesions and healthy control endometria. (A, B) Representative JUND staining shows increased expression in endometriosis lesions than control endometrium in both stromal cells and epithelial cells. (C, D) Representative CEBPD staining shows that CEBPD is expressed in both stroma and epithelium, and is highly expressed in the stroma of endometriosis lesion. E = epithelium, S = stroma.

## Discussion

It is known that endometrium is exposed to hypoxic periods specifically upon menstruation as well as during placentation (Pringle *et al.*, 2010; Maybin and Critchley, 2015). Here we conducted a global analysis of the transcriptomic responses to severe hypoxia in cultured ESF and DSC by comparing the hypoxia exposed (1% O_2_, 24h) cultures to normoxic controls. Overall our findings suggest that hypoxia establishes related but distinct transcriptional states in ESF and DSC. Several upregulated functional pathways, such as glycolysis and EMT, were shared with the two cell types, but many hypoxic responses, such as downregulation NFKB pathway in ESF and downregulation of stress responses as well as core hypoxic response genes in DSC were cell type specific. We also studied the relevance of these findings by intersecting our results with previous studies on endometriosis, endometriosis presenting a clinically relevant condition where stromal cells often inhabit hypoxic niches (Bishop, 1956; Bourdel *et al.*, 2007; Wu *et al.*, 2019), and investigated relevant discovered hypoxia regulated targets in the endometriosis and endometrium using immunohistochemistry.

Our results show that hypoxia interferes or modifies several gene regulatory networks necessary for early decidualization. In both ESF and DSC, hypoxia promotes fibroblastic phenotype (EMT), and thus, may interfere with the development of the quasi-epithelial state of DSC. Further, when looking at TFs, the intersection of our results with available PGR knockdown data (Mazur *et al.*, 2015) suggests that hypoxia partly reverses PGR dependent rewiring of transcriptional regulation typical for early decidualization.

In both ESF and DSC hypoxia enhanced classical hypoxia pathways including glycolysis and insulin-like growth factor (IGF) mediated signaling by altering the expression of IGF binding proteins (IGFBP1, IGFBP3 etc.). These modulate action of IGFs, and thereby may affect glucose uptake and proliferation of the cells (Ding and Wu, 2018). Substantial hypoxic induction of IGFBP3 has also been observed in other cell types (Natsuizaka *et al.*, 2012; Chang *et al.*, 2015) and has been associated with robust hypoxic polysome enrichment of IGFBP3 mRNA that may permit continuous translation under hypoxia (Natsuizaka *et al.*, 2012). In endometrial stromal cell increased glycolytic metabolism appears to take place also in well oxygenated conditions (Kommagani *et al.*, 2013), and the cells actively shuttle lactate (Zuo *et al.*, 2015). These observations can be also viewed in the context of Warburg effect or “pseudohypoxia” (Russell *et al.*, 2017), i.e. that stromal cells robustly express hypoxia-associated proteins, such as glycolytic enzymes regardless of oxygen levels. Moreover, recent studies on first trimester decidua show that the uterine natural killer cell (uNK) populations that are critical for placental development also drive glycolysis (Vento-Tormo *et al.*, 2018), suggesting that hypoxia (or pseudohypoxia) related metabolic programming including glycolysis is not limited to stromal cells and tumors but may be characteristic also to other cell types of decidualizing endometrium.

In our hypoxia experiment, stress responses appeared to be reduced specifically in DSC evidenced by general downregulation of stress response genes and the downregulation of mRNA levels of *HIF-1alpha* and *HIF-2alpha*, putatively representing negative feedback regulation via repressors such as upregulated *ID* genes (Inhibitor Of DNA Binding). Hypoxic downregulation of *HIF* mRNA in DSC can be viewed as part of general desensitized stress responses in DSC, as previously reported on cells exposed to oxidative stress (Kajihara *et al.*, 2013) and other stress pathways (Muter *et al.*, 2018) and may even be related to the above mentioned Warburg effect in DSC, as glycolysis short cuts the mitochondria which are major players in apoptosis. Transcriptional repression of *HIFs* do not contradict with the previously reported protein levels showing HIF2 stabilization during early secretory phase (Maybin *et al.*, 2018), because HIFs are mainly regulated by hypoxia on translational level (Bishop and Ratcliffe, 2014). Additionally, in our study, of the several upregulated repressor TFs, DECs (BHLHE40 and BHLHE41) specifically repress CLOCK and PER genes (Sato *et al.*, 2016), which were downregulated in our experiment. These circadian rhythm TFs are involved in synchronizing decidualization associated proliferation (Muter *et al.*, 2015), and our data suggest that also this switch is modified by hypoxia.

As early decidualization is partly an inflammatory process, several of the early decidualization factors are also regulators of inflammation, including STAT and CEBP pathways. In both cell types, but especially in DSC, hypoxia downregulated FOXO1, STAT3 and STAT5A that are known initiators of decidualization (Gellersen and Brosens, 2014), thus suggesting that hypoxia suppresses the initial pro-inflammatory phase of decidualization. Hypoxia also upregulated AP1 factors (JUND and JDP2) that often mediate inflammation, but that also are general activators for fibroblast proliferation (Manabe *et al.*, 2002; Florin *et al.*, 2004). On the other hand enhancers of decidualization *HOXA10, HOXA11, CEBPB* and *CEBPD* (Gellersen and Brosens, 2014) were upregulated by hypoxia, underscoring that hypoxic periods may also have positive effects for decidualization together with the potential to prime stromal cells for stress tolerance (Kajihara *et al.*, 2013) or glycolytic metabolism that is enhancing decidualization (Kommagani *et al.*, 2013; Zuo *et al.*, 2015).

Hypoxic modification of decidualization related inflammatory states or progesterone signaling may also be relevant in reproductive disorders including endometriosis, where hypoxia has been show to induce stromal cell migration and lesion growth (Lu *et al.*, 2014; Liu *et al.*, 2017). We investigated the relevance of our cell culture results by overlapping those with available transcriptomic data from endometriosis, and discovered a significant overlap of hypoxia upregulated genes with endometriosis upregulated genes. The overlap was most pronounced in a dataset of isolated endometrial stromal cells (Rekker *et al.*, 2017) in which endometriosis lesions were compared to healthy control endometrium. Notably, the high proportion of hypoxia upregulated genes (39%, 157/402) of the intersected endometriosis upregulated genes (Rekker *et al.*, 2017) suggests that hypoxic gene regulatory programs extensively contribute to the core gene regulation in the stroma of endometriotic lesions.

The overlap of 157 hypoxia and endometriotic stromal cell upregulated genes was enriched in genes promoting angiogenesis, EMT and estrogen responses. Angiogenesis and estrogen response were not among the most enriched pathways in the primary analysis of hypoxia upregulated genes (Fig. 1B) suggesting for the presence of endometriosis specific hypoxic interactions that are not present in stromal cells originating from healthy endometrium. Our results support previous observations (Wu *et al.*, 2012, 2019) that in endometriosis hypoxia and estrogen may synergistically promote the growth of the lesions. For example, the major angiogenic factor VEGFA was upregulated in our hypoxia experiment, and is known to be regulated by both estrogen and hypoxia (Zhang *et al.*, 2017). Among the angiogenesis enriched genes, we observed substantial hypoxic upregulation of KLF2, a known repressor of T-cells/monocyte activation and NFKB pathway (Jha and Das, 2017). Activation of NFKB pathway has been associated with the initiation of menstruation (Evans and Salamonsen, 2014), and in the context of endometriotic niche the downregulation of NFKB pathway may contribute to the ability of stromal cells to escape the clearance by immune system.

In order to systemically search for hypoxia upregulated TFs that drive endometriosis we conducted promoter analysis of the overlap of 157 hypoxia and endometriosis upregulated genes. We discovered that Jun/Fos family members JDP2 and JUND, and CEBP family members CEBPB and CEBPD were both robustly hypoxia upregulated, and enriched in the promoters of the overlapping genes. Of these, we selected JUND and CEBPD for protein level studies in deep endometriosis and healthy control endometrium and found elevated immunohistochemical staining of JUND and CEBPD in deep endometriosis compared to controls.

Our study highlights JunD as a potential hypoxia regulated intervention target in endometriosis. The upregulation of Jun/Fos pathway components in endometriosis has been reported in several studies (Hastings *et al.*, 2006; Tamaresis *et al.*, 2014) and previously C-JUN NH2-terminal kinase inhibitor was reported to reduce endometriosis in baboons (Hussein *et al.*, 2016). While cytokine profiling of endometriotic peritoneal aspirates detected Jun/Fos driven cytokine expression was interpreted as a major component of macrophage directed cytokine signature (Beste *et al.*, 2014), our results suggest that endometriotic stromal cells that express Jun/Fos factors contribute to the cytokine signatures of endometriosis.

*CEBPD*, which in our study was 5-fold induced by hypoxia in both ESF and DSC, has been suggested to act as a tumor suppressor that in hypoxia turns into a growth promoter (Balamurugan and Sterneck, 2013). Our immunohistochemical results showed increased CEBPD protein levels in deep endometriosis lesions. This differs from a previous immunohistochemical assessment of CEBPD in extra-ovarian endometriosis (Yang *et al.*, 2002), where no differential staining was reported between endometriosis and controls, potentially due to different types of lesions investigated. Further, CEBPD is a paralog of CEBPB, which is a known core regulator of decidualization (Wang *et al.*, 2012; Tamura *et al.*, 2017), however, both CEBPB and CEBPD target decidual PRL promoter (Pohnke *et al.*, 1999). Recent transcriptomic analysis have indicated that CEBPD is early upregulated and late downregulated during decidualization whereas CEBPB levels are not after the initial upregulation (Rytkönen *et al.*, 2019). Putatively, our observation of stronger hypoxia dependent transcriptomic upregulation of CEBPD together with the promoter analysis suggests that CEBPD may have more pronounced regulatory role in endometriosis compared to CEBPB.

Recently described HIF binding sites provide mechanistic perspective on hypoxia induced changes in the Jun/Fos or CEBP TF activities (Smythies *et al.*, 2019). For example, in HepG2 cells both HIF1 and HIF2 binding sites are enriched with CEBPD motifs (Smythies *et al.*, 2019), and in multiple cell types especially HIF2 binding sites are enriched with Jun/Fos motifs (Smythies *et al.*, 2019). Indirectly this suggests that also in endometrial stromal cells HIF2 is a potential interaction partner for Jun/Fos TFs. On the other hand hypoxia related regulatory interactions are not necessarily HIF mediated, but may proceed through changes in chromatin structure via oxygen dependent histone lysine demethylases (KDM). For example, ER receptor is regulated by KDM3A (Wade *et al.*, 2015), which is an oxygen dependent histone demethylase. Recently several KDMs, such as KDM5s (Batie *et al.*, 2019) and KDM6s (Chakraborty *et al.*, 2019), were reported to direct extensive hypoxia dependent transcriptional changes. Specifically, KDM6 activation has been associated with uterine fibroblast activation (Nancy *et al.*, 2018), which is concordant with our observation that hypoxia promotes fibroblastic phenotype in endometrial stromal cells.

In conclusion, the global transcriptome analysis performed in this study suggests that hypoxia enhances glycolytic energy metabolism and promotes fibroblastic phenotype (EMT) and, thus may, interfere with the development of the quasi-epithelial state of DSC. Hypoxia also modifies inflammatory pathways and partly counteracts PGR actions suggesting that hypoxia affects regulatory networks that are essential for decidualization. Hypoxia upregulated genes have significant overlap with previously detected endometriosis upregulated stromal genes (Rekker *et al.*, 2017) and promoter analysis of this overlap revealed that Jun/Fos and CEBP transcription factors are potential hypoxic drivers of transcription in endometriosis. Of these we validated the increased expression of JUND and CEBPD in endometriosis lesions using immunohistochemistry. Overall the findings suggest that hypoxic stress constitutes related but distinct transcriptional states in ESF and DSC, and that hypoxic regulatory programs contribute to the core gene regulation in the stroma of endometriotic lesions.

## Supplementary data

This is linked to the online version of the paper at XXX.

## Author contribution statement

KTR conceived and designed the study, conducted the cell cultures and RNA work, analyzed the data and wrote the manuscript. TH participated in the laboratory work and analysis. MM participated in the data analysis and writing of the manuscript. XM participated in the writing of the manuscript. AP contributed in the sampling and participated in the writing. LLE provided resources for the work and participated in the writing. MP and GPW participated in the design of the work, writing of the manuscript and provided resources for the work.

## Acknowledgements

We are grateful to Roger Babbitt and Jordan Pober lab (Yale University) for the help with hypoxia cell culture. We thank Erica Nyman and Sinikka Collanus from the University of Turku Histology core facility and Satu Orasniemi from the Institute of Biomedicine for their contribution in immunohistochemistry.

