## Supplementary Figures for "Transcriptomic responses to hypoxia in endometrial and decidual stromal cells"

**Supplementary Figures 1 and 2:**

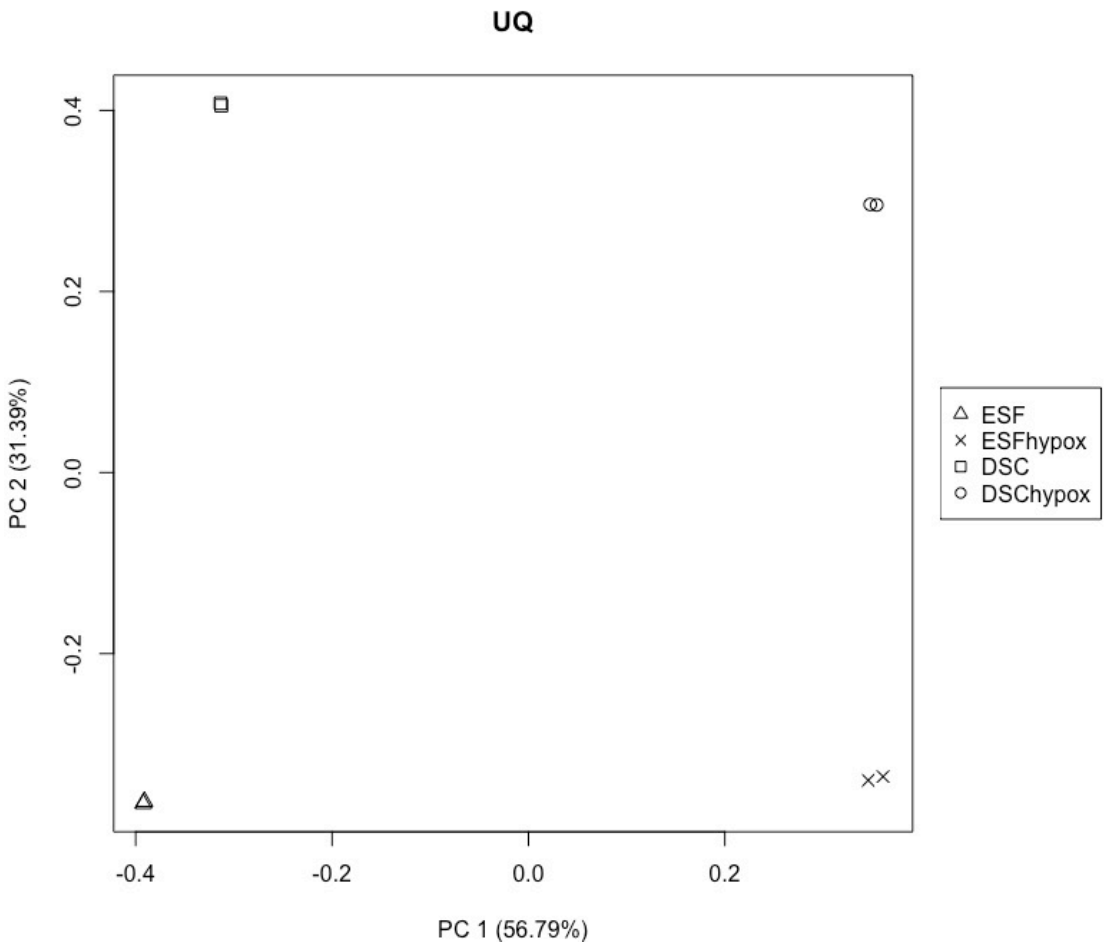


**Supplementary Figure 1.** The PCA analysis of the four conditions (ESF, DSC, ESF hypoxia and DSC hypoxia) after edgeR upper quartile normalization displays that the two biological replicates inside each experimental condition group tightly together. PC1 separates hypoxia and normoxia conditions and PC2 separates differentiation state (ESF or DSC).

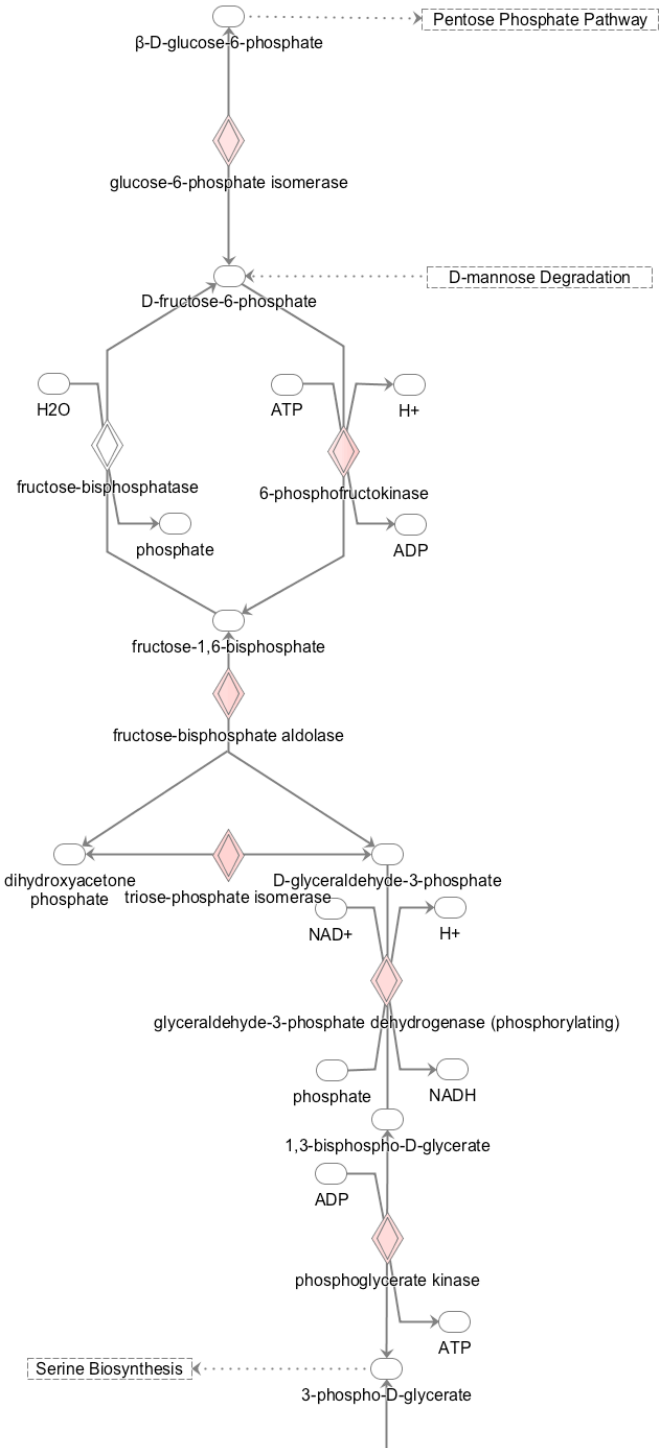


**Supplementary Figure 2.** Subset of Glycolysis pathway showing hypoxia upregulated enzymes in both ESF and DSC (in red, FDR < 0.01, FC > 2.0, TPM > 2).
